# Anatomical tool in the identification of ploids in maize seedlings and potential use in initial stage of double haploides obtainment process

**DOI:** 10.1101/777292

**Authors:** Raquel M. O. Pires, Genaina A. Souza, Danielle R. Vilela, Heloisa O. Santos, Renato C. C.Vasconcellos, Édila V. R. Von Pinho

## Abstract

Studies that optimize the haploid technique in the removal of maize lines are necessary. Between the stages that mostly requires attention and it is directly related to the success of the technology is the correctly separation of induced haploids and diploids. Morphological markers are commonly used but have strong influence of the environment, and laboratory methods have been developed and may be more efficient. Thus, the objective was to study the use of the anatomical analysis tool, through the analysis of young maize leaf for use as the indirect markers in the identification of ploidys. The hybrids were crossed with the KEMS haploid inducer. The seeds crossed, were selected according to the R-navajo marker and submitted to two different protocols of chromosome duplication. Plants that survived to the duplication protocols were acclimated in greenhouse and then transferred to the field. After the self-polinization of the DH0 plants, the DH1 seeds were taken to the field, divided into treatments according to the parentals and duplication protocols. At the vegetative stage V4 of the plants, leaf tissue samples were collected to the evaluation of the amount of DNA and identification of ploidys and anatomical analysis. The nuclear DNA review of each sample was performed for the comparison in histograms of the position of G1 peak to the G1 peak of the internal or external reference standard. A high accuracy came to validate an anatomical tool, through the variables studied in this work, as a marker in the differentiation of ploidis in maize plants, and it can be used in selection programs. The anatomy made in some letters is a non-destructible technique and, together with a flow cytometry technique, can be used as an indirect method in haploid cutting programs at the initial stage of the identification of seedlings.

## Introduction

The success of a breeding program that aims at the production of maize commercial hybrids lies in the fact of obtaining elite lines. Among all the steps, this is considered to be the most time-consuming and costly, and the technology of double haploid emerges as a way of reducing time in obtaining these lines [1].

The rapid production of homozygotic lines allows a better exploitation of genetic variability and increases the efficiency of selection. Homozygous plants will have the maximum additive variance, the effects of dominance and epistasis, and the advantages in the selection of quantitative, superior characteristics [2]. In addition, the decrease of the costs with labor, use of smaller experimental area and anticipation of profits in commercial programs for maze breeding have made this technique a great success.

The production of double-haploid lines involves four main steps: in vivo induction of haploidy, identification of possible haploids, chromosome doubling and the self-fertilization of lines obtained for increment of seeds [3]. However, the success of this methodology is still dependent on the use of inductors with high capacity of induction, a precise system of identification and differentiation of haploid and diploid seeds, as well as efficient and reproducible protocols of chromosome doubling [4].

The doubling of the chromosome number spontaneously or induced by the application of mitotic agents e.g., colchicine, retrieves the diploid condition and restores fertility [5]. The action mechanism of colchicine involves the irreversible connection to tubulin dimers, causing a conformational change and preventing the polymerization of mitotic spindle, and as a result, the newly duplicated chromosomes are not separate and the core reorganizes with the number of duplicated chromosomes [6]. However, not all cells of the treated tissue polyploidize, which can lead to the formation of chimeras, i.e., tissues or plants with duplicate sectors and others unduplicated ones [7] called mixoploids. Truly duplicated lines resulting from this process are called duplicate or double haploids (DHs).

There are several methods for certification of polyploidization, being flow cytometry the most used one [8]. Flow cytometry is a reliable and fast method, because it allows the analysis of a large number of cells and of different tissues [9]. In experiments with double haploids, flow cytometry allows vigorous seedlings and detected as diploids in the histograms are discarded before the step of field, reducing time and space. In addition, flow cytometry allows the analysis of the efficiency of the protocol of chromosome doubling, since it is not possible to confirm if the response of seedlings to duplication was positive [10–12].

Another tool that is being studied as a marker in the differentiation of ploidies in plants is the leaf anatomy, being the leaf considered the component with greater ability to adapt to environmental conditions. Highly flexible, leaf anatomy is influenced by environmental factors, such as, irradiation (leaves of sunlight/shade, [13], nutrients [14], drought [15] and ozone [16,17]. Changes in the leaves characteristics, such as those related to the thickness of the leaf blade, parenchymas, epidermis and number of stomata for example, and that are highly associated with the photosynthetic potential of plants, are used in studies of genetic selection by the use of morphological and anatomical markers. Additionally, they are highly heritable characteristics, i.e., can be passed to their offspring [18].

The cytoanatomic characterization is a methodology that allows the identification of haploid and supposed polyploidy in plants subjected to chromosome doubling. The study of measurement and comparison of stomata, based on the principle that the length of the same normally increases with the number of chromosomes, is the most commonly cited in the literature [19].

The number of stomata in association with other leaf anatomical characteristics, has already been mapped to different levels of ploidy in studies with coffee plants [20]. For this species, the greater the number of stomata the higher the ploidy. In the case of *Coffea canephora*, a reduction in the stomatal density is higher in the tetraploid level for some cultivars [20]. [21], observed in Citrus that the size and density of stomata varied according to ploidy level, where the triploids showed a higher number of stomata when compared to diploid plants. A similar result was observed by [22], who stated to be possible the use of anatomical markers for purposes of selection of citrus with different levels of ploidy.

Ploidy is well studied from the point of view of genetics and genomic perspective, but the morphological and anatomical aspects related to these differences in the amount of DNA, remain poorly studied in maze plants. Analyzing the anatomical characteristics of young leaves of maize, capable of discriminating the different ploidies and extrapolate these results in diploid and haploid discrimination on the optimization of the process of obtaining double haploid is of extreme importance.

Thus, the objective of this work was to study the use of anatomical tool, through the analysis of the characteristics of young leaves of maize for use as indirect markers in the identification of ploidies, and through future studies, to extrapolate the use of this marker in the identification of haploids in the initial stage of the process of obtaining double haploids.

## Material and methods

The seeds used in this work were obtained from an experiment previously developed by [23] through the cross between four simple hybrids (DKB393, GNS 3225, GNS 3264, GNS 3032) with the haploid inducer KEMS, used as male parental. Seeds from these crosses were separated by staining of the embryo and endosperm and selected as possible haploids according with the marker R-navajo [24].

The authors submitted the haploid seeds to two chromosome duplication protocols, and the plants that survived the field, called DH0, that produced pollen and had stigma style in synchronism, were self-fertilized, resulting in DH1 generation.

Thus, in this present work, in order to evaluate the maintenance of DH in future generations, the DH1 ears were harvested and the seeds threshed and dried at room temperature up to 12% moisture level at the point of physiological maturity. The seeds were then mixed and divided into treatments as shown in Table 1. These seeds were then stored in a cold chamber at 10 ° C until the following experiments were carried out.

**Table 1.** Treatments taken for the field in the harvest season 2014/2015, established in accordance with the parental and the protocols of chromosome doubling.

| Materials Identification |  |  |
| --- | --- | --- |
| Treatments | Hybrids | Protocols |
| 1 | DKB393 | 1 |
| 2 | DKB393 | 2 |
| 3 | GNS3225 | 1 |
| 4 | GNS3225 | 2 |
| 5 | GNS3264 | 1 |
| 6 | GNS3264 | 2 |
| 7 | GNS3032 | 1 |
| 8 | GNS3032 | 2 |

In the harvest season 2014/2015, the total number of seeds DH1 for each treatment, was taken to the field and the experimental design was a randomized complete blocks with eight treatments with four replications. Each block was composed by lines of 10 meters of length, with spacing of 80 cm between rows and among plants of 25 cm, and sowing of a seed per hole. The collection of leaf material occurred in young stage of the plant maize, with 4 replicates collected at random within each treatment.

In the case of leaf anatomy, 5 replicates were performed for each repetition in the laboratory, five blades of 10 sections each, were made, and the top five fields were photographed and subsequently measured. The following were measured: thickness of the leaf blade (ELF), thickness of the parenchyma (PAR), thickness of the upper epidermis (EES) and lower epidermis (EEI), polar diameter of the stomata in the upper face (DPS) and lower (DPI), equatorial diameter of the stomata of superior face (DES) and lower face (DEI) and stomatal density on the upper surface (DS) and lower (DI). For that, in the vegetative stage V4 which corresponds to the development of four leaves, leaf tissue samples were collected.

For that, in the vegetative stage V4 which corresponds to the development of four leaves, leaf tissue samples were collected.

For both evaluations, the medial portion of the young leaves fully developed, was cut into segments of approximately 3 cm in length and wrapped in aluminum paper duly identified per plant. Aluminum papers remained in a polystyrene box containing recyclable crushed ice, until the time of transportation to the laboratory of tissue culture of plants of the Department of Agriculture of UFLA.

At the opening of each envelope of aluminum, part of the samples of leaf tissue was used for the quantification of DNA and part to the anatomical analysis. Therefore, young leaf tissue samples of maize were crushed under ice, in Petri plates containing 1 mL of cold buffer LB01, to obtain nuclear suspension [25], which was added 2,5µL RNase and stained with 25 µL of propidium iodide (1 mg mL-1). The species *Vicia faba* (quantity of DNA of 26.9 pg/2C) was used as an external standard of reference and for each sample at least 10 thousand cores were analyzed. Each suspension was analyzed in flow cytometry FacsCalibur (Becton Dickinson). The histograms obtained were evaluated by the WinMDI software 2.8 (2009) for the evaluation of the peaks of DNA. The estimate of the nuclear DNA content (pg) of each sample was performed by comparing the position of the peak G1 with the peak G1 of the internal standard or external reference.

For this comparison the following expression was used:

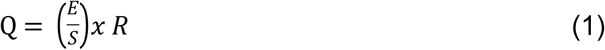

Where:

Q is the quantity of DNA of the evaluated sample (pg/2C).

E is the position of the G1 peak of the sample.

S is the position of the peak G1 reference standard and

R is the quantity of DNA of the standard sample (26.9 pg/2C).

By the quantity of DNA it was possible to make inferences about the ploidy level of the evaluated genotypes.

As mentioned, the other part of the samples of leaf tissue was immediately fixed in FAA_50_, (formaldehyde: acetic acid: ethanol, 5:5:90) for 48 h. Then, the samples were removed from the fixative solution, rinsed and stored in 70% ethanol solution [26]. At this moment the samples were transported to the Laboratory of Anatomy and morphogenesis of Plant Biology Department of the Federal University of Viçosa, where permanent slides were made using portions of the leaf that were dehydrated in ethyl series included in historresin (methacrylate), according to the manufacturer’s recommendations. Transverse sections of leaf were sectioned in automatic advance rotary microtome (model RM2155, Leica Microsystems Inc., Deerfield, USA) with 5 μm thick, arranged on histological slides and stained with toluidine blue [27], to the limbal micro morphometry.

For the stomata evaluations, fragments of the central part of the leaf blade were sectioned from material stored in alcohol 70%. The samples were clarified by means of the technique of Diaphanization, described by [28] and modified for the species, having been clarified in methanol for 48 hours and then in lactic acid for 6 hours in water bath for 98 °C and assembled into lactic acid. After the staining procedure, the samples were immersed in ethanol 80, 70 and 50%, and later were rinsed in distilled water. The histological slides with leaf fragments were mounted on glycerol-jelly Images of the slides of cross cuts and diaphanization were obtained under a light microscope (model AX-70 TRF, Olympus Optical, Tokyo, Japan) coupled to a digital camera (model Zeiss AxioCam HRc, Göttinger, Germany) and a microcomputer with the program to capture images Axion Vision, having been digitized and stored in a microcomputer. For the analysis, 10 distinct fields of each sample were measured by means of Image-Pro^®^ Plus software (version 4.1, Media Cybernetics, Inc., Silver Spring, USA).

The analyzes were performed for all characters with the estimation of variance components and the prediction of random effects using the approach of mixed models, the method of restricted maximum likelihood/best linear unbiased prediction (REML/BLUP) [29]. For this, the following statistical model was used:

Where:

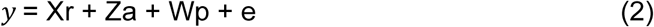

*y*: vector of data;

R: vector of the effects of repetition (assumed as fixed) added to the overall average;

A: vector of genotypic effects among treatments (random), being 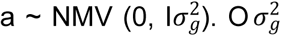 is the variance associated with genotypic among treatments;

p: vector of genotypic effects among treatments (random), being 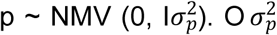 is the variance associated with the parcels effects;

e: vector of random errors, and 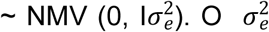 is the variance associated with the residual effects;

X, Z and W: incidence matrices for r, and p, respectively.

The heritability in the average of treatments 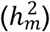 and accuracy were estimated, and the significance of the random effects as the genotypic variance among treatments 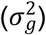 were tested by the *Likelihood Ratio Tests* (LRT) where the analyses of deviance were obtained for each character evaluated [30].

These statistical analyzes were performed using the software SELEGEN-REML/BLUP [29].

## Results

For all the evaluated characteristics the accuracy was classified as very high [31], with values above 87.0%. On average, the heritability 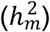 of the characteristics was also high with values above 84.0%, what indicates good reliability of data (Table 2). The heritabilities of treatments ranged from 76.0% (EES) to 98.0% (DEI) among the evaluated characteristics, which demonstrates that in the anatomic evaluation in this experiment, the largest part of the variation observed is due to genetic causes to the detriment of environmental variations [32].

**Table 2.**
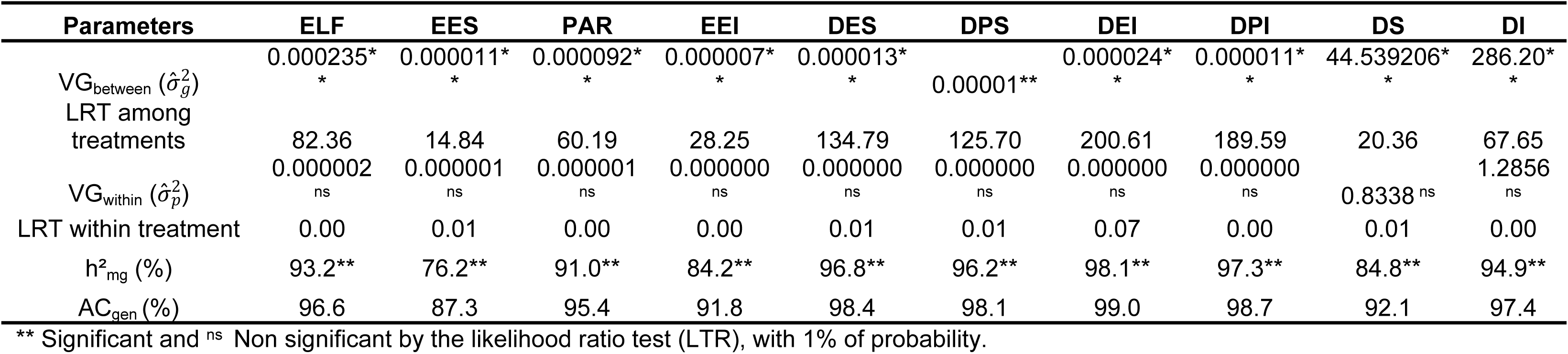
Summary of ANADEV and estimates of genetic parameters in the evaluation of maze hybrids for traits. Thickness of the leaf blade (ELF), thickness of the upper epidermis (ESS), thickness of the parenchyma (PAR), thickness of the lower epidermis (EEI), equatorial diameter of the stomata in the upper face (DES), polar diameter of the stomata in the upper face (DPS), Equatorial surface of the stomata in the lower face (DEI), polar diameter of the stomata of the lower phase (DPI) and stomata densities of upper surfaces (DS) and lower (DI) of the leaf.

The genotypic variance was highly significant by the LRT test and likelihood ratio for the characters related to the thickness of the leaf blade (ELF), stomatal density of both the upper surface (DS), as well as the lower surface (DI), thickness of the parenchyma (PAR), thickness of the upper epidermis (EES) and lower epidermis (EEI), polar and equatorial diameter of upper epidermis (DES and DPS) and lower (DEI and DPI), according to LRT (P < 0.01), characterizing these characteristics as good candidates to be inserted in a breeding program as markers. It was possible to realize that the characteristics related to the thickness of the lamina and those related to the stomata were the ones that showed higher heritability, followed by the thickness of the parenchyma. The thickness of the epidermis, even with high heritability, were lower (Table 2).

The anatomical differences observed were obtained through anatomical cuts and diaphanization of samples. It was possible to observe a difference in leaf blade as the thickness (Fig 1) and quantity and size of the stomata (Fig 2).

**Fig 1.**
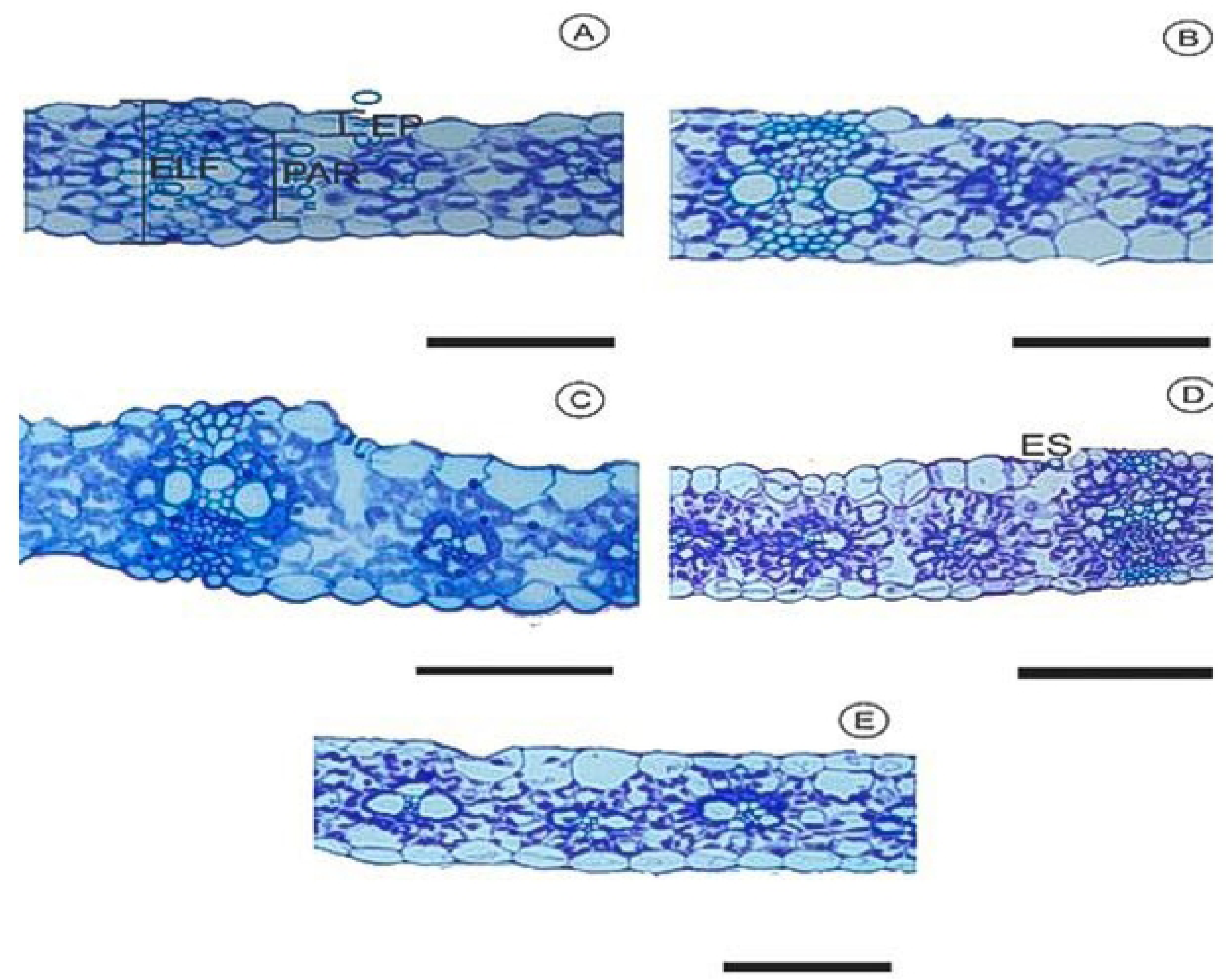
Cross-sectional cuts of maze leaves collected after a process of chromosome doubling with use of colchicine. Haploid material (A), diploid (B), Double haploid (C), triploid (D) and tetraploid (E). ELF= thickness of leaf blade, PAR= thickness of parenchyma, EP=epidermis and ES= stomata. The bars represent 50 µm.

**Fig 2.**
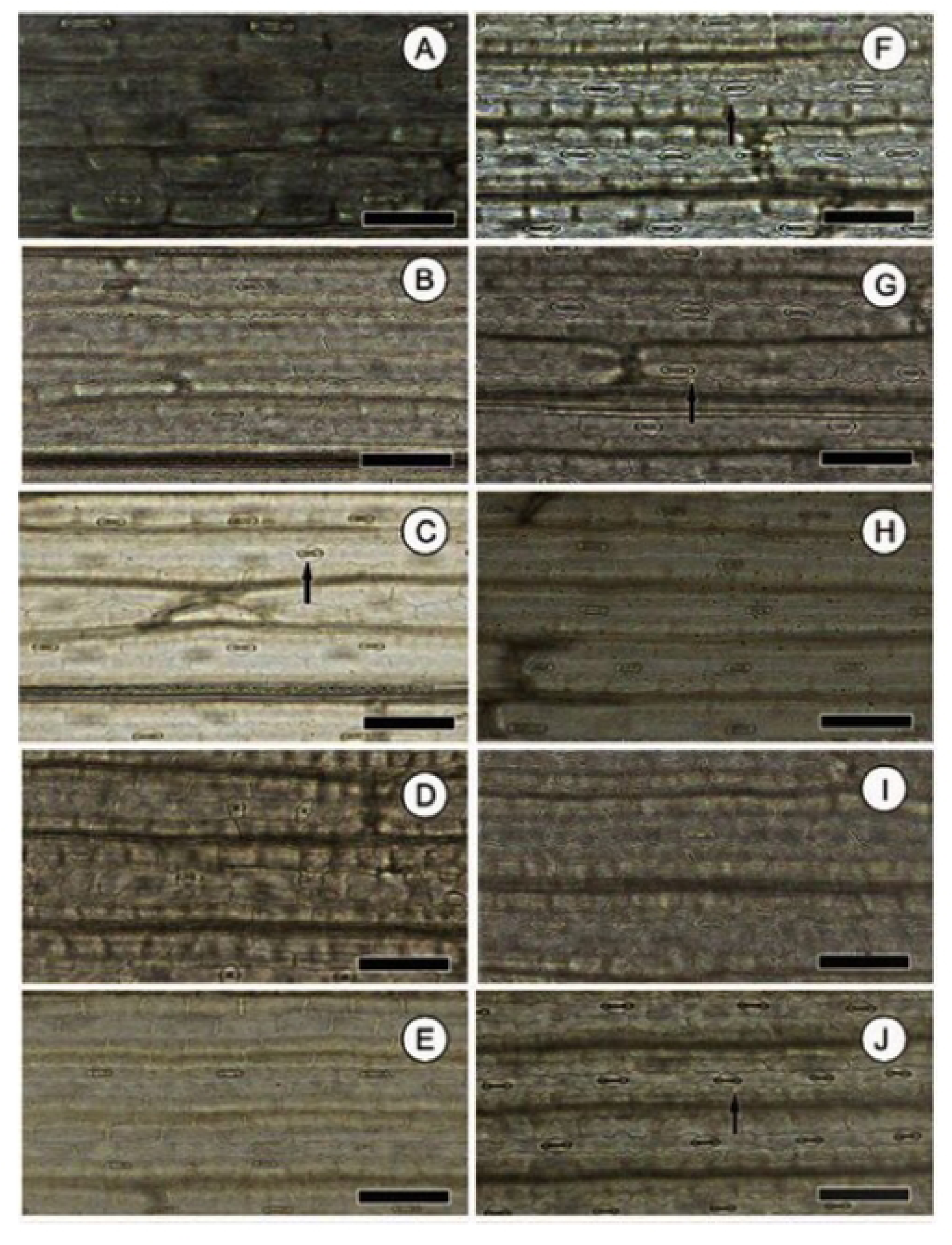
Adaxial surface (A to E) and abaxial (F through J) of maze leaves collected after a process of chromosome doubling with use of colchicine. Haploid material (A and F), diploid (B and G), Double haploid (C and H), triploid (D and I) and tetraploid (E and J). The bars represent 50 µm and arrows indicate stomata.

In Table 3, it is observed that the treatment 7 presented the highest values of ELF, ESS, EEI and held until the third position in the ranking for PR, DEI, DS. While the treatment 6 occupied until the second position to ELF, ESS, PR, DPS and DPI. The treatments 1 and 5 occupied the last two positions to ELF, EES, PR, EEI, DES, DPS, DEI and DPI.

**Table 3.**
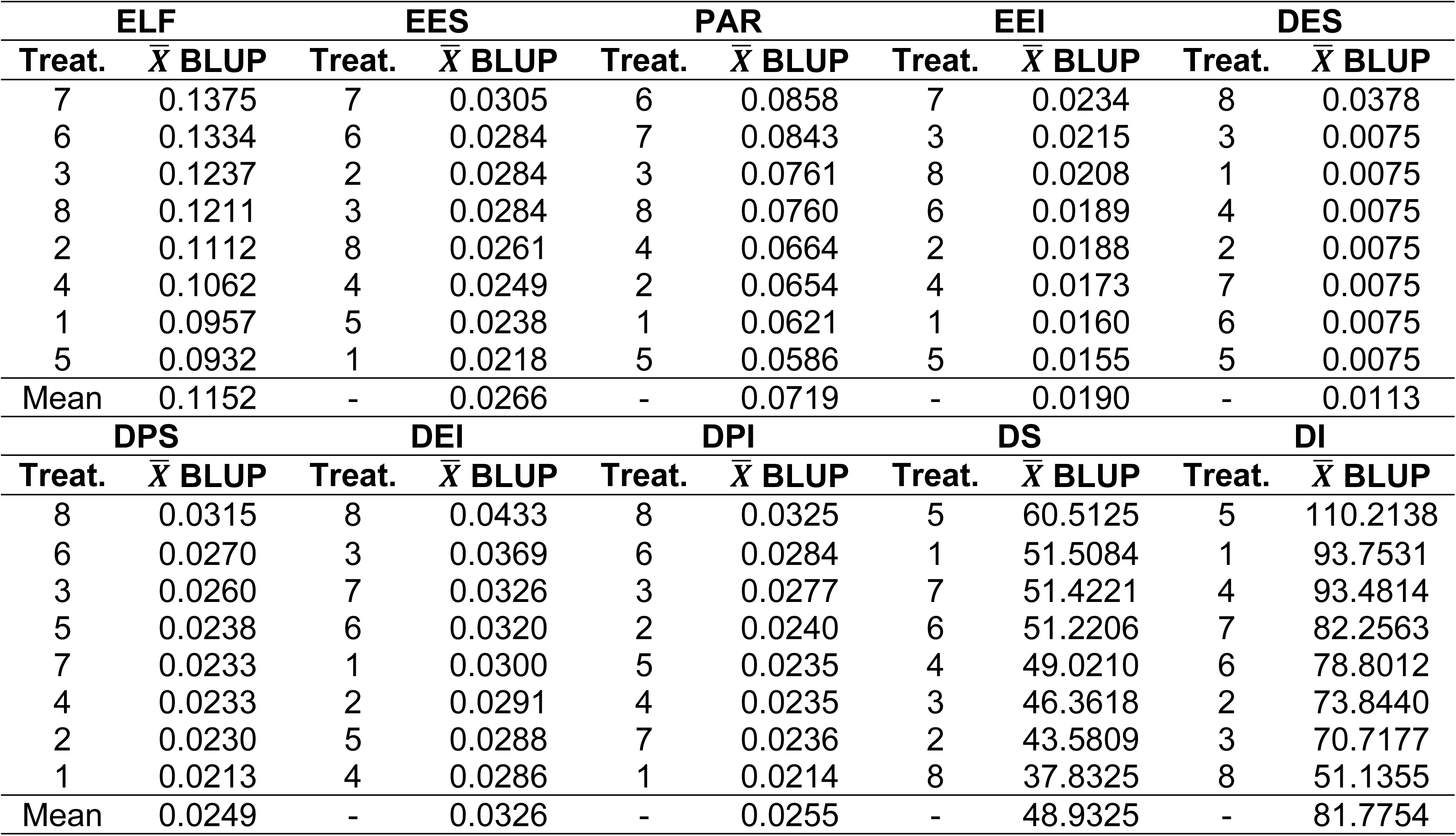
Genotypic averages and ranking of treatments for the anatomical characteristics related to leaf blade. Leaf blade thickness (ELF), thickness of the upper epidermis (ESS), thickness of the parenchyma (PAR), thickness of the lower epidermis (EEI), equatorial diameter of the stomata in the upper face (DES), polar diameter of the stomata in the upper face (DPS), polar and equatorial diameter of the stomata in the lower face (DEI) and (DPI), stomatal density of the upper surfaces (DS) and lower (DI) of maze leaves collected after a process of chromosome doubling with use of colchicine.

The importance or contribution of each analyzed variable as a possible marker in the separation of the tested plants, demonstrates greater relevance of the thickness of the leaf blade and the stomata, while the thickness of epidermis little contributed to the separation of plants. (Fig 3).

**Fig 3.**
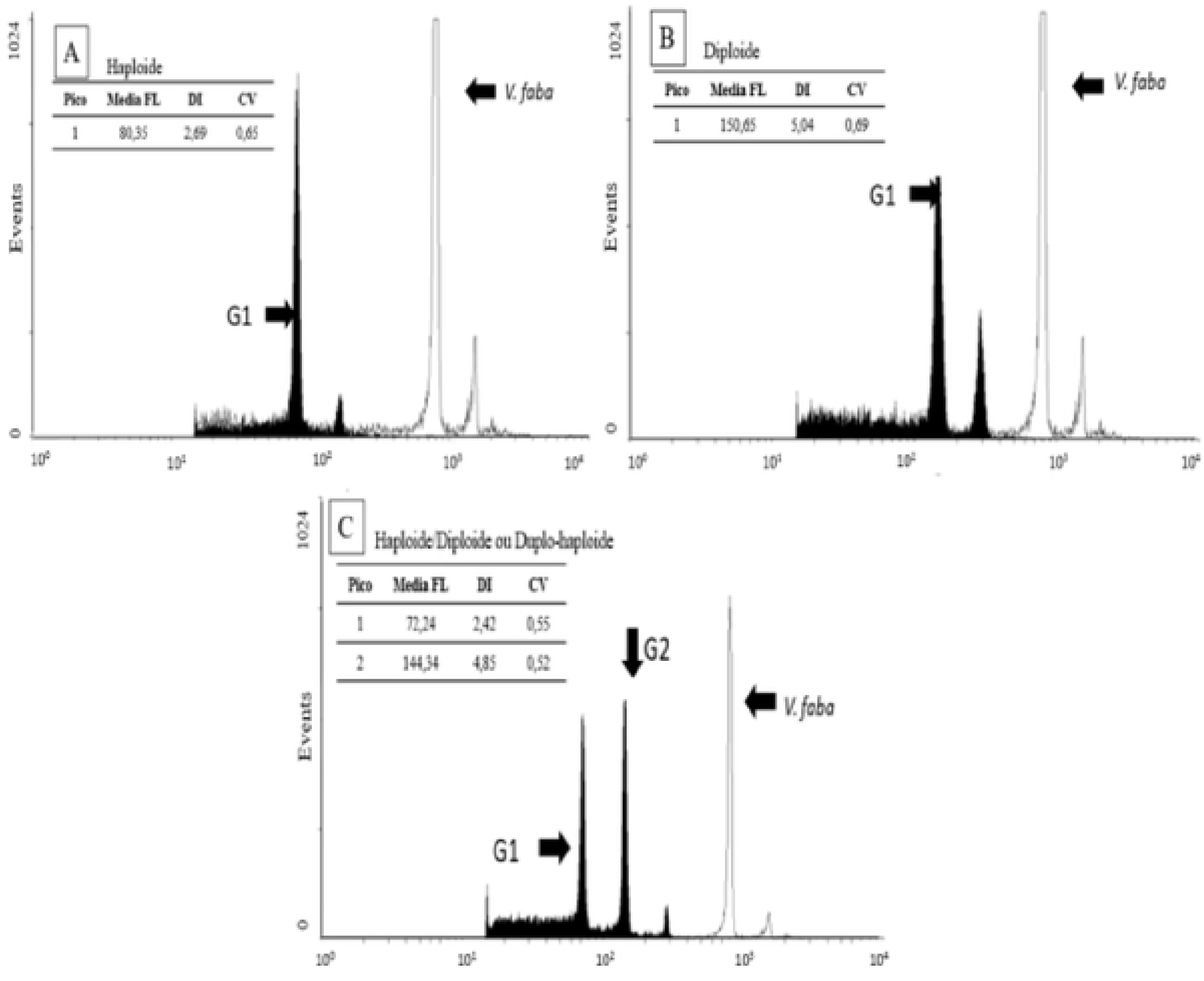
The relative importance of the characteristics evaluated in the separation of the valuated materials.

In parallel to the anatomical analyzes and with the aim of inferring the real ploidies observed in the tested materials, the DNA quantification was performed by flow cytometry technique in 32 selected plants (Table 4). The histograms obtained by this method allows for the identification of the ploidy level of the individuals tested through the location of the G1 peak of the sample on the axis of the relative intensity of fluorescence (Fig 4). The dominant peaks generated in the histograms are relative to the quantity of DNA of the cores in the G1 phase of the cell cycle. The estimate of the ploidy level is done by comparing the G1 peaks of the histogram of a sample with the peak of a plant-standard with known ploidy [25].

**Table 4.**
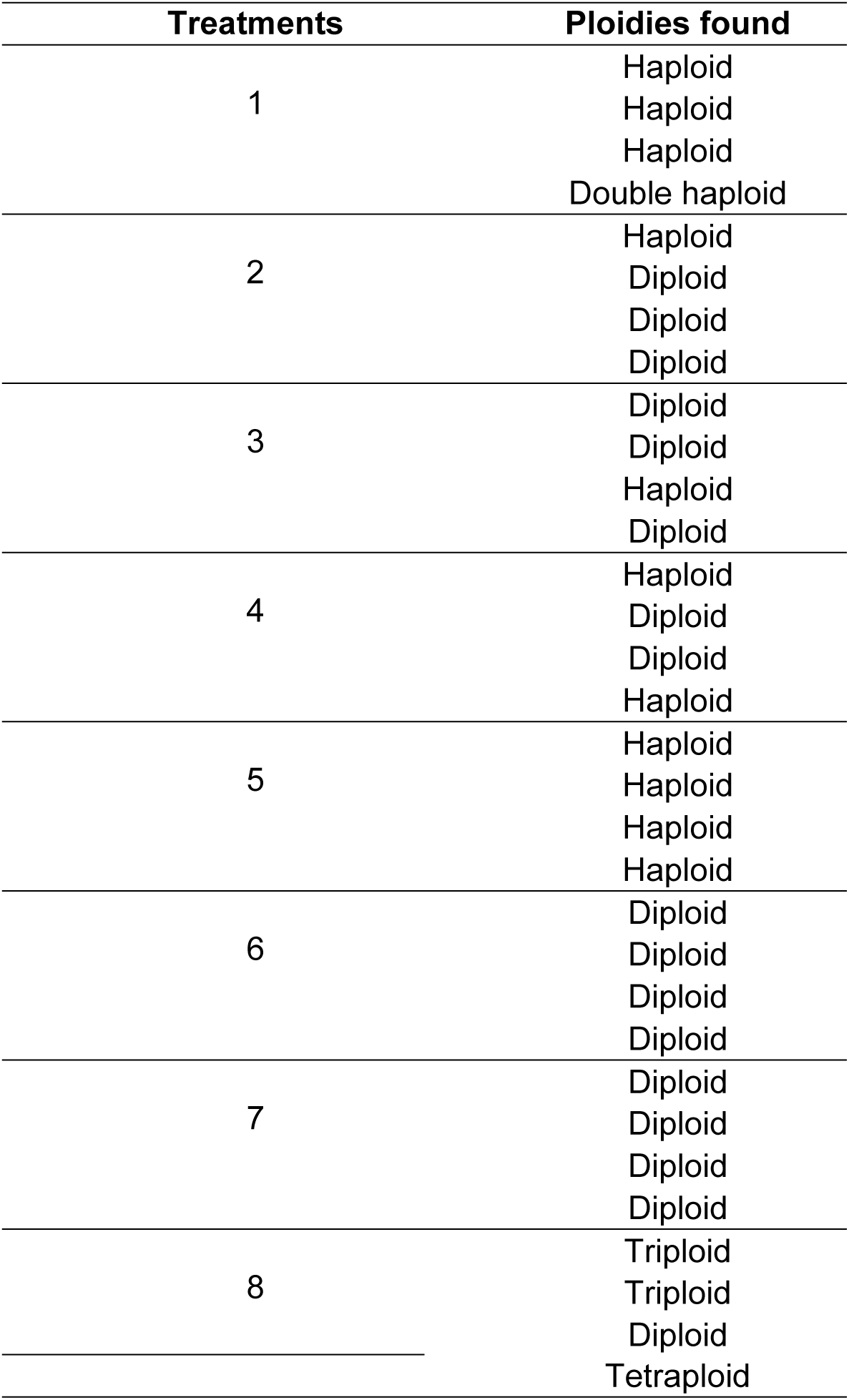
Identification of the ploidy level in agreement with the analysis of flow cytometry and flow of 32 plants evaluated divided by treatment.

Maize diploids have the peak G1, located in the region of relative intensity of fluorescence, soon after the mark of 10^2^ (Fig 4A). The haploids have lower relative intensity of fluorescence and the peak G1 is located to the left of the mark of 10^2^, i.e., dislocated in the direction of the x axis (Fig 4B). Whereas the triploids and the tetraploids have still greater intensity than the diploids, locating at the right of the mark of 10^2^ and with higher peaks (fig not shown). Whereas the Double haploid plants, are those that have ploidies haploid/diploid type (Fig 4C).

Usually, an association of the histogram obtained with a histogram of a standard plan is performed, for example, *Vicia faba*, used by [10,11,33]. The use of standard allows to obtain the quantity of DNA contained in the sample.

**Fig 4. Histograms of ploidies detected by flow cytometry in maize plants collected after a process of chromosome doubling with use of colchicine. A. Diploid Plant B. Haploid Plant. C. Double haploid Plant**. Vertical axis = number of read cores; horizontal axis = relative intensity of fluorescence. The arrows show the peaks G1 and G2 and the external standard of reference.

## Discussion

It was possible through the analysis of mixed model to verify the existence of variability in the anatomical characteristics, i.e., there is a difference among the tested materials (P < 0.01) (Table 1). In addition, these variables showed a quality required for insertion in a breeding program, with the aim of separation of evaluated materials [34]. The heritability and accuracy obtained were high in accordance with the classification made by [31] (Table 1). The fact of the evaluated anatomical characteristics have potential for use in selection programs is of extreme importance, since these characteristics may be associated with the photosynthetic potential of plants, i.e., the productive capacity of plant material, leaves, roots and seeds.

The thickness of the leaf blade (ELF), has a crucial role not only in the capacity of carbon fixation by the chloroplasts of the palisade parenchyma, but also by the internal storage of CO_2_ by sponge parenchyma [35]. While the stomata are the channels of influence of CO and the flow of water vapor. For the plants to be effective, they must balance the gaseous exchanges carried out through these structures to maximize the absorption of CO_2_ for photosynthesis and minimize the loss of water through transpiration. Thus increasing the efficiency of the use of the water and consequently the plasticity of the plant in the face of environmental changes. A program that aims at obtaining hybrids with greater adaptive capacity, seems a major bottleneck of agriculture through the global climate changes [36].

The stomata behavior, therefore, controls the volume of CO_2_ in the intercellular spaces of the leaf for photosynthesis. Even if the maze as plant of C4 metabolism is able through the mechanisms of CO_2_ concentration, to maintain an adequate quantity of C for photosynthesis [37], the stomatal density and the size of the stomata are important characteristics to maximize efficiency. Once that, in spite of the area of the pores of the stomata represent less than 3% of the total area of the leaf, about 98% of all the absorbed CO_2_ and water lost occurs by these pores [38].

The anatomical characteristics of the stomata define the stomatal conductance (*gs*), theoretical maximum [39], i.e., the functionality of the same and also influence the speed of response. The *maximum gs* relates to the size and density of stomata, which can be influenced by the environment of growth [40,41]. However, as in this study, all the plants were grown in the same environment, we can consider that the density and the pattern of size, influenced by the atmospheric concentration of CO_2_, water availability [42] and light [43], varied according to the genetic characteristics of each tested hybrid. This reinforces the importance of the anatomical characters as early markers of separation of hybrids used in this study.

Experimental evidences showed that the density of stomata is negatively correlated with the stomata size [40,41]. The interaction/correlation among stomata size and density, and the impact on stomatal function has received much attention, particularly with reference to the evolution of the performance and plasticity in plants [41]. Evidence from several studies have also suggested that smaller stomata respond faster than larger stomata, an observation that has been explained in the context of relations surface-volume and the requirement for ck to boost the movement [44].

The selection of plants grown with changes in the density of stomata to increase the performance of plants has been widely exploited [38,45], with limited success. The increase of the stomatal density can increase the *gs* and the photosynthetic rate can become 30% greater in conditions of high brightness [46].

The increase of photosynthesis can encourage the increase in weight, as already mentioned. Increase in the weight of seed has also been associated with induced polyploidy [47]. What can potentiate the vigor and germination of seeds, favoring the formation of a more homogeneous stand. However, the manipulation of functional stomate responses is clearly more complicated, requiring a thorough understanding of the metabolism of the manipulated plant.

The use of these anatomical characteristics of the leaf is widely used for identifying the levels of ploidy in many species of plants, such as alfalfa [48], *Gossypium* [49], *Dactylis* [50], ryegrass [51], wheat [52] and *Bromus inermis* [53,54]. In coffee the density of stomata decreased while its size increased with an increase in the ploidy level, with the lower density found in the tetraploids and higher in the diploids [20]. Genotypic differences in stomatal frequency and length of the guard cells were also observed in barley [55], soybean [56] Triticale [57,58]. These studies demonstrate the possibility of the use of anatomical markers with mechanism for identification of ploidy.

In the present, it is verified the contribution of the ten variables evaluated in separate studied plants and coincides with what is reported in the literature in relation to the great importance of the stomata. Carefully observing the relative contribution of each trait, it is verified that the variables associated to the stomata represent, altogether, approximately 50% of the contribution of separation (Fig 3).

The high contribution of leaf blade is due to its constitution. The leaf blade is composed by parenchyma, in which chloroplasts and spaces for CO_2_ storage are located, in addition of course, all the other components of the leaf. Therefore its relevance is easily understandable and the importance of variables related to the stomata is also evident, reinforcing what is already described in several academic articles.

In addition it is possible to suggest that the anatomic variables, as possible markers were efficient on grouping even partially the hybrids (Tables 2 and 3). Behavior that reinforces what has been described above, where the traits were efficient anatomical markers for various crops. The treatments that showed higher averages for the analyzed variables were classified as diploids (Table 3), in general with averages exceeding the haploids (Table 2).

The increase in ploidy level may be the main driving force to facilitate the plants breeding, as it provides important phenotypic effects, such as increasing the size of the cells and organs, and sometimes a larger force and biomass, and additional molecular and phenotypic variation that may arise soon after the formation of the polyploids. This behavior can be attributed to the effect “gigantism”, in which plants with higher ploidy may have increased the size of their structures [19]. The treatments 6 and 7 (diploids), for the characteristics related to leaf blade (ELF and PAR), showed higher averages to the haploids (Table 2). However the effect “gigas” was not observed to hybrids tri and tetraploids, as reported in *bulbophyllum ipanemense* [59].

Significant effects on the ploidy level, and the anatomical and morphological characteristics, such as leaf dry mass and thickness of the epidermis, have already been reported in Brassicas [60] and characteristics such as leaf thickness and photosynthetic rate, for rice [61]. The increase in the leaves thickness and total mass of plants may result in greater energy expenditure, however, as the maze is a plant of Kranz anatomy, there is not so much spent on histodifferentiation of juxtaposed layers of palisade parenchyma, since these cells are found around the cells of the sheath of the beam. Therefore the gain in leaf thickness would contribute not only to the increase of the total mass of the leaf, but also to the increase of empty spaces. These spaces play an important role in the CO_2_ reserve for photosynthesis and because it does not require energy to histodifferentiation, being less costly in terms of energy.

The size of the cells and the thickness of the components were positively correlated with the ploidy level also in potatoes [62]. With the increase of the genome, the gigantism in cells and organs is widely observed and associated with the increments in the photosynthetic rate [63]. The increase of photosynthesis is attributed to the increase of the tissues, increased capacity for storage of CO_2_, and increase of *gs*. The size of the epidermal cells, cells, can also be associated with the ploidy level of the material under observation [47]. In addition to the increase in the activity of multiple enzymes such as hydrolases and expansins [64].

The effect “giga” was also related in previous studies with modifications of cell wall through a loosening, which enables a higher rate of growth of plants and phenotypic changes. This loosening is assigned to a higher expression of genes of expansin enzyme in rice [65,66], tobacco [67,68] and Arabidopsis [69,70]. The role of expansins would be to induce the extent of cell wall, generating larger cells, higher plants and longer roots. So in these cases, the cell expansion associated with the ploidy is related to the increase of molecular signaling for synthesis of genes of expansin and may lead to an increase in weight of structures, as in tomato [71].

## Conclusions

The thickness of the leaf blade and the size of the stomata are highly heritable traits in maize.

The obtained high accuracy validates the anatomical tool through the variables studied in the present work, as a marker in the differentiation of ploidies in maize plants, which may be employed in programs for selection of hybrids.

The anatomy made in young leaves of maze is a non-destructible technique and in conjunction with the technique of flow cytometry, can be used as indirect method in programs to obtain double haploids, in the initial stage of identification of seedlings.

## Author Contributions

**Conceptualization:** RMOP HOS EVRVP

**Formal analysis:** RMOP GAS

**Funding acquisition:** EVRVP

**Investigation:** RMOP GAS DRV RCCV

**Writing:** RMOP DRV RCCV

